# Off-target glycans encountered along the synthetic biology route towards humanized N-glycans in *Pichia pastoris*

**DOI:** 10.1101/2020.01.28.923631

**Authors:** Bram Laukens, Pieter P. Jacobs, Katelijne Geysens, Jose Martins, Charlot De Wachter, Paul Ameloot, Willy Morelle, Jurgen Haustraete, Jean-Christophe Renauld, Bart Samyn, Roland Contreras, Simon Devos, Nico Callewaert

## Abstract

**Background:** The glycosylation pathways of several eukaryotic protein expression hosts are being engineered to enable the production of therapeutic glycoproteins with humanized application-customized glycan structures. In several expression hosts, this has been quite successful, but one caveat is that the new N-glycan structures inadvertently might be substrates for one or more of the multitude of endogenous glycosyltransferases in such heterologous background. This then results in the formation of novel, undesired glycan structures, which often remain insufficiently characterized.

**Results:** When expressing mouse interleukin-22 (mIL-22) in a *Pichia pastoris* (syn. *Komagataella phaffi*) GlycoSwitchM5 strain which had been optimized to produce Man_5_GlcNAc_2_ N-glycans, glycan profiling revealed two major species: Man_5_GlcNAc_2_ and an unexpected, partially α-mannosidase-resistant structure. A detailed structural analysis using exoglycosidase sequencing, mass spectrometry, linkage analysis and NMR, revealed that this novel glycan was Man_5_GlcNAc_2_ modified with a Glcα-1,2-Manβ-1,2-Manβ-1,3-Glcα-1,3-R tetra-saccharide. Also the biosynthetic intermediates of this off-target modification were detected. Expression of a Golgi-targeted GlcNAc Transferase-I strongly inhibited the formation of this novel modification, resulting in more homogeneous modification with the targeted GlcNAcMan_5_GlcNAc_2_ structure. We have also observed the off-target glycan on other glycoproteins produced in the GlycoSwitchM5 strain. This illustrates the intricacies of Golgi glycosylation pathways and cautions that the use of glyco-engineered expression host cells should always be accompanied by detailed glycan analysis of the particular therapeutic proteins being produced.

**Conclusions:** Our findings reinforce accumulating evidence that robustly customizing the N-glycosylation pathway in *Pichia pastoris* to produce particular human-type structures is still an incompletely solved synthetic biology challenge, which will require further innovation to enable safe glycoprotein pharmaceutical production.

## BACKGROUND

Glycosylation is the most widespread posttranslational modification of secreted and membrane-associated eukaryotic proteins. N-glycosylation is required for the efficient folding of many glycoproteins through the recruitment of calnexin/calreticulin-associated ERp57 protein disulfide isomerase (1). For proteins that require such glycan-assisted folding, *E. coli* is unsuitable as an expression system and eukaryotic host cells are then used instead. However, different eukaryotic cells modify the N-glycans in various ways in the Golgi apparatus. For instance, yeasts generally produce high/hyper-mannose N-glycans that largely comprise of mannose, whereas mammalian cells produce mainly complex-type N-glycans, in which the glycan branches are modified with N-acetylglucosamine (GlcNAc), galactose (Gal), fucose (Fuc) and sialic acid (Sia). However, the pharmacokinetic and pharmacodynamic behavior of therapeutic glycoproteins is strongly influenced by the type of the glycans they carry. For example, yeast-produced high-mannose N-glycan modified glycoproteins are cleared from the bloodstream very rapidly through binding to (hepatic) macrophage lectins with mannose specificity (2). This is disadvantageous if long circulatory half-life is desired, but advantageous if delivery to the macrophage endosomal system is intended, as is for example the case in Enzyme Replacement Therapy of Gaucher disease. Nevertheless, yeast’s native glycans can be immunogenic, and even when a high-mannose glycan is desired on the therapeutic glycoprotein, the yeast pathway needs to be significantly engineered to produce a human-type high-mannose structure, such as Man_5_GlcNAc_2_. Indeed, much work is being invested in such re-engineering of glycosylation pathways in eukaryotic expression host cells (3). For fungal hosts such as *Pichia pastoris*, the common theme in most of these pathway engineering efforts consists of two parts. First comes the inactivation of the main host-cell specific glycosyltransferases which steer the host-specific pathway away from a glycan precursor structure that is common with the human pathway, resulting in the accumulation of this precursor. Most often, this is Man_8_GlcNAc_2_. Second, heterologous glycosidases and glycosyltransferases are introduced, which modify the common precursor to the desired structures (often mammalian-type high-mannose, hybrid or complex N-glycans). For instance, to reach the Man_5_GlcNAc_2_ structure, which is a high-affinity ligand for the macrophage mannose receptor, an α-1,2-mannosidase is targeted to the ER-Golgi boundary that converts Man_8_GlcNAc_2_ into Man_5_GlcNAc_2_. In previous work, we have named the resulting strain as ‘GlycoSwitchM5’. The Man_5_GlcNAc_2_ glycan is also the starting substrate for the human N-glycosylation pathway, which can subsequently be introduced (4).

Such N-glycan engineering has been thoroughly explored in *Pichia pastoris*, in our group and formerly at Glycofi Inc. (USA). Indeed, research demonstrated that strains can be obtained that produce human-type N-glycans. Nonetheless, the published data only report on a limited number of test-case proteins, and in most cases the glycan analytics were focused on the major glycan species, disregarding low abundant components in the mixture. The production of larger protein quantities (developed for true pharmaceutical use) enabled deeper analysis that often leads to the detection of unexpected glycoforms. Indeed, dozens of the host cell’s endogenous glycosyltransferases remain active in the Golgi apparatus of the engineered cells (5). For example, it has been described that glyco-engineered *P. pastoris* strains can incorporate β-1,2-linked mannose residues in some of their N-glycans (6), a modification believed to be only present on the outer chain of mannoproteins and O-glycans of *wild-type Pichia*-derived glycoproteins (7, 8). Reportedly, such structure has also been found on a human monoclonal IgG produced in glyco-engineered *P. pastoris* (9). The problem could be largely circumvented by identifying and then knocking-out the likely glycosyltransferase culprits. Nevertheless, it is clear that careful analysis remains necessary for each glycoprotein expressed in glyco-engineered strains.

Here the knowledge on this understudied aspect of glyco-engineering is extended by presenting the structural analysis of another unexpected N-glycan found on *Pichia* GlycoSwitchM5-produced interleukin-22 (IL-22), a member of the IL-10 superfamily of cytokines (10, 11, 12). In the GlycoSwitchM5 strain, Man_5_GlcNAc_2_ typically makes up >80 % of all N-glycans on proteins produced (13). However, this was decidedly different in the case of mIL-22, as reported in this study, offering an opportunity to perform detailed structural analysis of the off-target glycans.

## RESULTS

### Characterization of *Pichia* GlycoSwitchM5-produced mIL22

Interleukin-22 has three N-glycosylation consensus sites and all three can be modified with an N-glycan (14). Murine IL-22 expressed in the wild-type GS115 strain appears as four major molecular species on SDS-PAGE (Figure 1, A), ranging in molecular weight from approximately 17 to >40 kDa. The upper bands (bands 2, 3 and 4) disappear after PNGase F endoglycosidase digestion, which indicates that these are N-glycoforms of IL-22, whereas the lower band (band 1) corresponds to the non-N-glycosylated protein.

**Figure 1.**
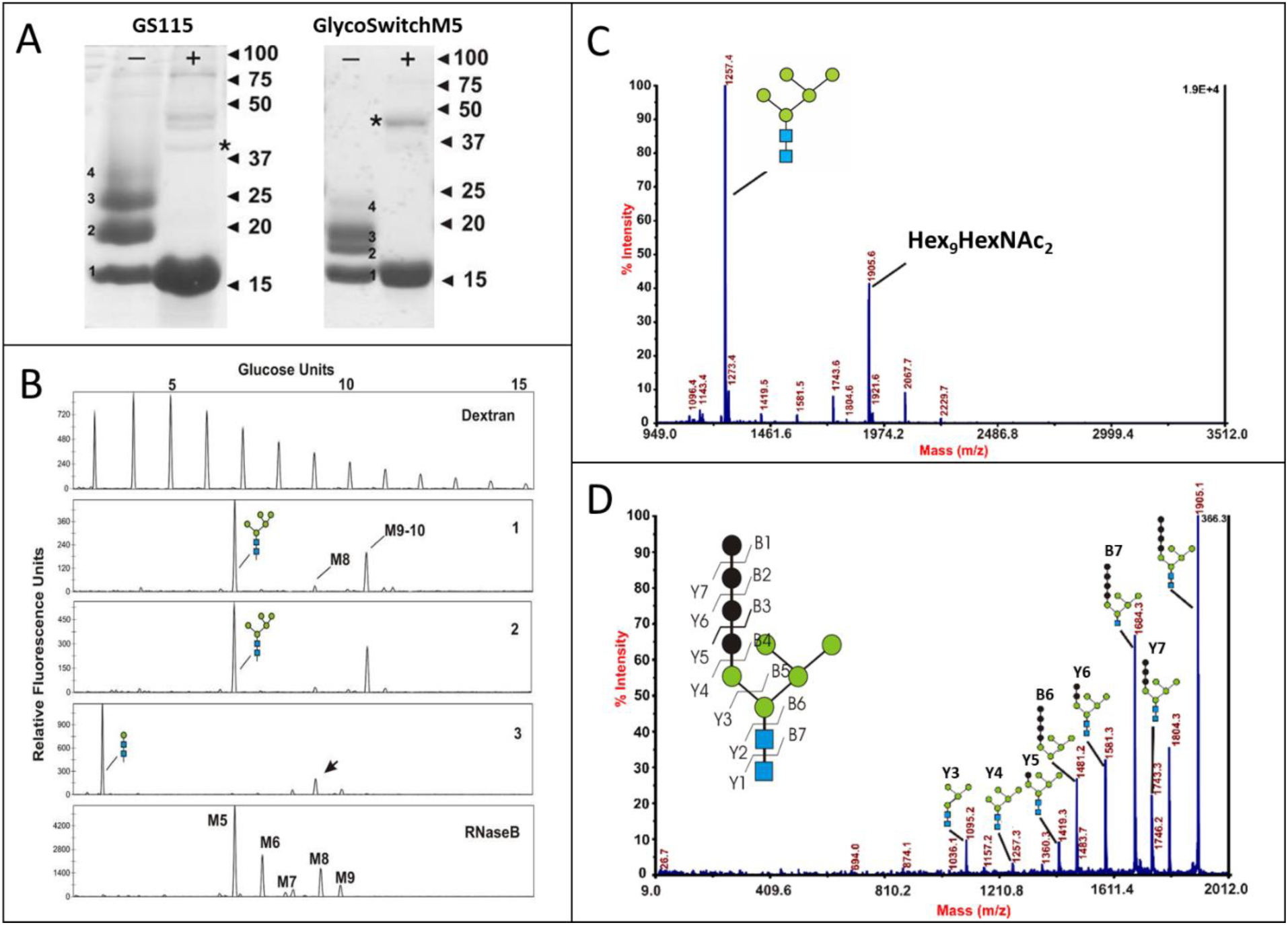
Characterization of GlycoSwitchM5-produced mIL-22 glycoforms. **A.** SDS-PAGE analysis of GS115- and GlycoSwitchM5-produced mIL-22, untreated (-) or treated with PNGaseF (+). Four glycoforms are distinguishable in the untreated sample: 1 = no N-glycans; 2 = 1 N-glycan; 3 = 2 N-glycans; 4 = 3 N-glycans. The band marked with an asterisk is PNGase F. **B.** DSA-FACE analysis of PNGaseF-released N-glycans. Top: Dextran reference ladder; Lane 1: untreated mIL-22 GlycoSwitchM5 N-glycans; Lane 2: α-1,2 mannosidase digested N-glycans; Lane 3: Jack Bean α-1,2/-3/-6-mannosidase digested N-glycans; standard N-glycan mix from RNAseB. **C.** MALDI-TOF MS analysis of released N-Glycans. **D.** MS/MS fragmentation of the Hex_9_HexNAc_2_ species (m/z 1905.6).

In parallel, a GlycoSwitchM5-mIL-22 expression strain was generated. Similar to wild-type produced mIL-22, purified GlycoSwitchM5-mIL-22 appears on SDS-PAGE as four major bands, ranging in molecular weight from 17 to about 25 kDa (Figure 1, A), all consistent with the modification of mIL-22 with a less heterogeneous (and lower molecular weight) N-glycan composition, as intended in this strain. Enzymatic removal of the N-glycans with PNGase F again resulted in a single non-N-glycosylated species. The bioactivity of mIL-22 was tested in a Ba/F3m22R cell proliferation assay and this showed that GlycoSwitchM5-produced mIL-22 is at least as active as mIL-22 produced in *E. coli* (specific activity of 7.9 U/µg for *E. coli* produced mIL-22 compared to 14.33 U/µg for GlycoSwitchM5 produced mIL-22).

### Structural analysis of N-glycans of purified Man_5_GlcNAc_2_-mIL-22

After PNGase F release, the N-glycans were labeled and analyzed by fluorophore-assisted carbohydrate capillary electrophoresis on a DNA sequencer (DSA-FACE) (15) (Figure 1, B). The electrophoresis profile showed that in addition to the expected Man_5_GlcNAc_2_ peak, a second major species migrated as a Man_9-10_GlcNAc_2_ N-glycan in the electrophoresis profile (Figure 1, B; lane 1). In order to gain more insight into the structure of this putative N-glycan, exoglycosidase digestions were performed. When digesting the N-glycans with α-1,2-mannosidase no shift was observed (Figure 1, B; lane 2). In contrast, the unknown N-glycan species had an increased mobility by about 1.5 glucose units upon Jack Bean α-mannosidase digestion (Figure 1, B; lane 3, arrow), consistent with the loss of two α-mannosyl residues (compared to the difference in mobility of the M7 and M9 reference N-glycans of RNaseB). The Man_5_GlcNAc_2_ N-glycan was completely digested down to the Man_1_GlcNAc_2_ N-glycan core after Jack Bean α-mannosidase digestion (Figure 1, B; lane 3), indicating that the Jack Bean mannosidase digest was complete. The apparently altered relative peak area of the two structures after Jack Bean digestion is due to the more efficient electro-kinetic injection of the very small tri-saccharide, which is inherent to this analytical method. The loss of two α-mannosyl residues upon broad specificity α-mannoside digestion is consistent with the hypothesis that the novel structure is likely a derivative of the Man_5_GlcNAc_2_ N-glycan in which one of the 3 terminal α-mannoses is modified.

Next, the PNGaseF-released N-glycans were analyzed by MALDI-TOF MS (Figure 1, C). In agreement with the DSA-FACE analysis, two dominant peaks appear in the spectrum: one peak corresponds to Hex_5_GlcNAc_2_ (Na^+^ adduct; m/z 1257.4) and the other to Hex_9_GlcNAc_2_ (Na^+^ adduct; m/z 1905.6). Also minor signals are found that correspond to Hex_8_GlcNAc_2_ and Hex_10_GlcNAc_2_. Fragmentation of the Hex_9_GlcNAc_2_ (m/z 1905.6) by MALDI-TOF/TOF MS/MS provided evidence of a linear tetra-hexoside modification on the Man_5_GlcNAc_2_ core N-glycan (Figure 1, D). This confirms that there is indeed only a single, linear modification on one of the terminal branches of the Man_5_GlcNAc_2_ N-glycan. To quantify the relative abundance of the different N-glycans from mIL-22 more accurately, an MS analysis was performed after permethylation of the N-glycans (Supplementary Figure 1, A). These results show that roughly ∼20% of the sample consist of a Hex_9_GlcNAc_2_-type N-glycan relative to the Man_5_GlcNAc_2_ N-glycan (Supplementary Figure 1, B). Moreover, a series of lower-abundance glycans were present with increments of one hexose residue up to Hex_10_GlcNAc_2_.

Finally, through GC-MS linkage analysis, the monosaccharide content of the N-glycan mixture and their linkage was determined (Supplementary Table 1). To this end, the permethylated N-glycans were acid hydrolyzed and the resulting sugar residues were analyzed by GC-MS. In addition to the expected linkages of mannose and GlcNAc in the main Man_5_GlcNAc_2_ N-glycan, also terminal glucose, 3-linked glucose, 2-linked mannose and 3-linked mannose was detected, with an approximated relative abundance ratio of 1:1:2:1. This is consistent with the CE and MS data that showed an abundance of the modified N-glycan of approximately 20-25% relative to the Man_5_GlcNAc_2_ N-glycan. Either one of the two 2-linked mannoses or the 3-linked mannose must be the mannose residue of the Man_5_GlcNAc_2_ structure that carries the linear tetra-hexoside substituent. Moreover, the data showed that a glucose residue is found at the terminus of this substituent. The presence of glucose residues in this glycan was particularly surprising, as it has never been described in N-glycans of mature, secreted proteins in this species.

### NMR-spectroscopy

To further elucidate the structure of the substituent, an NMR analysis was performed on the PNGaseF released N-glycans, to reveal the sequence and the anomericity of the glycosidic linkages. First, a 1D ^_1_^H-spectrum at 700 MHz was acquired. The analysis was performed on the mixture of N-glycans, as this assists in assigning the resonances belonging to the substituent based on the relative peak intensities. Upfield in the 1D ^_1_^H-spectrum, the signals that correspond to the methyl groups of the acetyl-function of the 2 GlcNAc-residues can be distinguished (1.8 – 2.2 ppm) (Supplementary Figure 2). In the anomeric region, 4 anomeric signals were eliminated from further consideration based on their very low intensities relative to the other anomeric signals (Figure 2, A; top trace). These very low intensity anomeric signals likely come from a small percentage of native *Pichia* N-glycans, containing 2-linked and 2,6-linked mannose, as seen in the linkage analysis. Furthermore, diffusion analysis was used to eliminate two more signals that do not belong to any of the N-glycan species but originate from smaller saccharide constituents present in the sample, possibly degradation products (Supplementary Table 2, marked with * and **). Exclusion of these contaminants leaves 12 signals (A-L).

**Figure 2.**
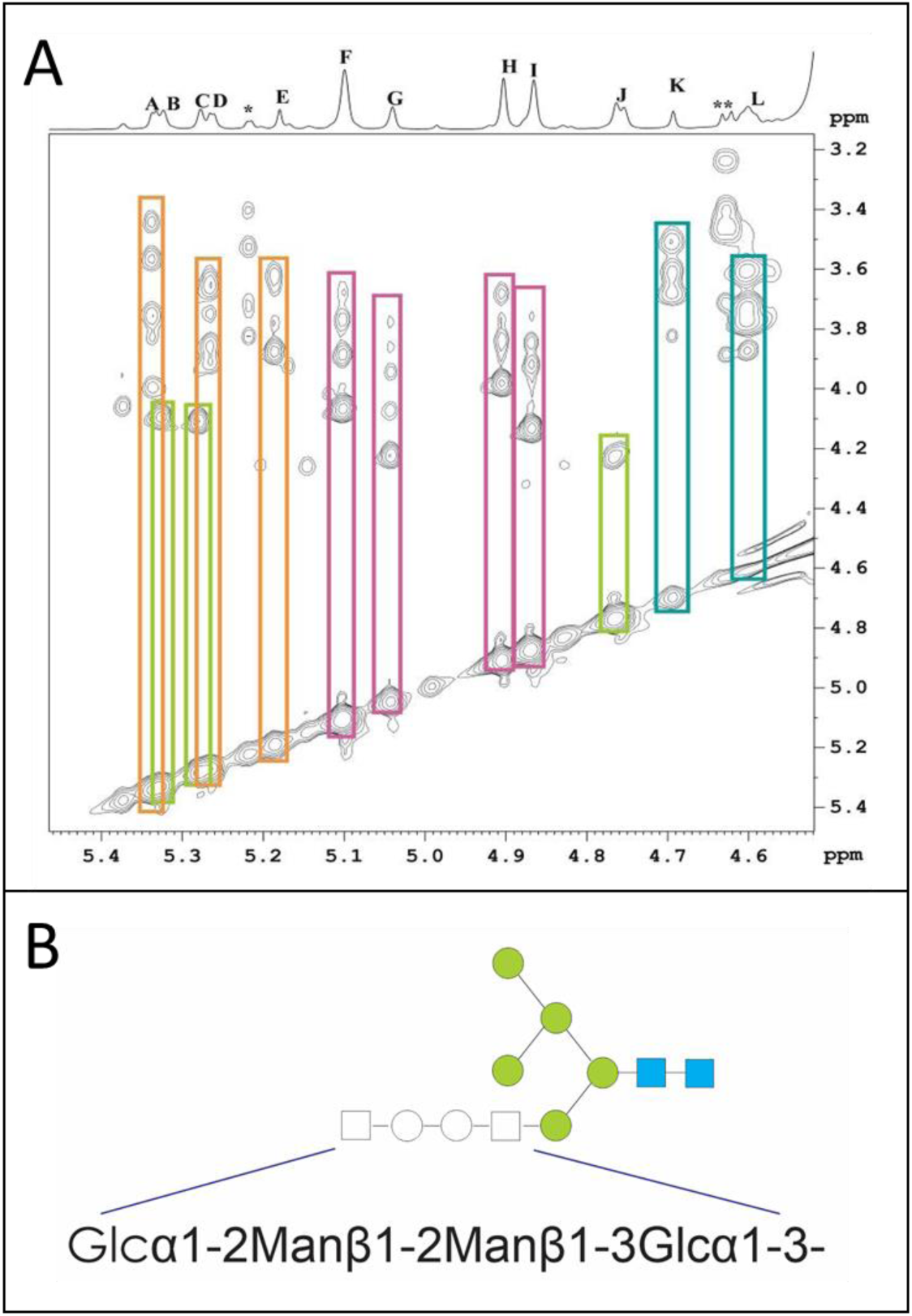
Structure determination of the unexpected N-glycan from GlycoSwitchM5-produced mIL-22 with NMR. **A.** 1D ^1^H NMR spectrum and TOCSY spectrum of the unexpected Hex_9_GlcNAc_2_ N-glycan. TOCSY matching allows to assign the monosaccharide ring geometry and anomericity: orange (A, D, E): α-D-glucose-type; green (B, C, J): β-D-mannose-type; pink (F, G, H, I): α-D-mannose-type; blue (K, L): β-D-glucose-type. **B.** Proposed structure of the unexpected N-glycan based on the NMR data and the MALDI- and GC-MS data.

Then, Total Correlation Spectroscopy (TOCSY) was used to identify the monosaccharide constituents of the tetra-saccharide substituent on the Man_5_GlcNAc_2_ N-glycan based on the 12 remaining anomeric signals (Figure 2, A). The specific stereochemistry of different monosaccharides causes a distinct TOCSY-trace for each, which allows for a relatively easy identification (16). The TOCSY traces of B, C and J (Figure 2, A; green traces) have a single crosspeak, which is characteristic for the β-D-mannose stereochemistry. Based on the integrated peak intensity, residue J is the β-mannose in the core of the N-glycans, which is common to all, whereas B and C have a lower peak intensity, consistent with these being part of the Hex_9_GlcNAc_2_ glycan. Thus, both 2-linked mannose residues in the tetra-saccharide substituent are β-linked. A typical TOCSY pattern for monosaccharides of D-glucose geometry has three or more distinct crosspeaks of similar intensity and the crosspeaks tend to be very broad (because of overlapping ringproton resonances). Such distinct traces are found for A, D and E, as well as for K and L. Based on the difference in their chemical shift, traces A, D and E were assigned as having α-D-glucose-type stereochemistry (δ(H1) > 4.8 ppm) (Figure 2, A; orange traces) and traces K and L as β-D-glucose-type stereochemistry (δ(H1) < 4.8 ppm) (Figure 2, A; blue traces). The remaining TOCSY traces F, G, H and I all have a pronounced crosspeak followed by a series of less distinct crosspeaks, which is characteristic for α-D-mannose stereochemistry (Figure 2, A; pink traces). The common Man_5_GlcNAc_2_ core-structure consists of 4 α-D-mannoses and a single β-D-mannose, in addition to one β-GlcNAc (with β-glucose ring geometry) and the reducing terminus GlcNAc which can be both the α- or β-epimer. Based on the high intensities of the anomeric resonances: F, G, H and I originate from the 4 α-mannose residues in the common core, J is from the β-mannose residue, L is from the β-GlcNAc, and H is from the β-anomer of the reducing-end GlcNAc. One of either A, D or E is from the α-anomer of the reducing-end GlcNAc. This allows to conclude that the four monosaccharides that constitute the tetra-hexoside are a terminal α-glucose, two 2-linked β-mannoses and a 3-linked α-glucose.

Heteronuclear Single Quantum Coherence (HSQC) spectroscopy indeed confirmed the monosaccharide content of the tetrasaccharide, and it also allowed to determine the type of linkages within the tetrasaccharide (Supplementary Table 3). One glucose residue was placed on the most distal end in an α-linkage, whereas the other 3-substituted glucose residue is α-1,3-connected to a terminal mannose of the Man_5_GlcNAc_2_ core. In between both glucose residues, there are two β-linked mannose residues, bound on the anomeric carbon (C1) and at their C2 position. This set of constraints leaves only 1 possible structure, i.e. Glcα-1,2-Manβ-1,2-Manβ-1,3-Glcα-1,3-Manα1-R (Figure 2, B).

### Characterization of GlycoSwitchM5-hIL-22

Murine IL-22 and human IL-22 have a high amino acid sequence similarity (>80%). Therefore, it was investigated whether GlycoSwitchM5-produced hIL-22 carries similar N-glycans as found on mIL-22. G115- and GlycoSwitchM5-produced hIL-22 leads to a comparable SDS-PAGE pattern as compared to mIL-22 (data not shown), and Man_5_GlcNAc_2_-hIL-22 is also functionally active (specific activity of 6.5 U/µg for *E. coli* produced hIL-22 compared to 19.37 U/µg for GlycoSwitchM5 produced hIL-22).

The GlycoSwitchM5-hIL-22 was purified, and its N-glycan composition was analyzed by DSA-FACE, as described above. This revealed a second major peak in addition to the expected Man_5_GlcNAc_2_ peak, however this N-glycan was situated around 9 glucose units. A lower abundant peak was also present just above 10 glucose units, quite similar to the one found on GlycoSwitchM5-mIL-22 (Figure 3; lane 1) (it should be noted that some relative shifts in the DSA-FACE electropherograms can occur, as these were not yet internally calibrated and the measurements on mIL-22 and hIL-22 were performed several years apart). Comparable to mIL-22, the 9-glucose units peak was recalcitrant to α-1,2-mannosidase (Figure 3; lane 2) and was partially digested after Jack Bean mannosidase treatment (Figure 3; lane 3).

**Figure 3.**
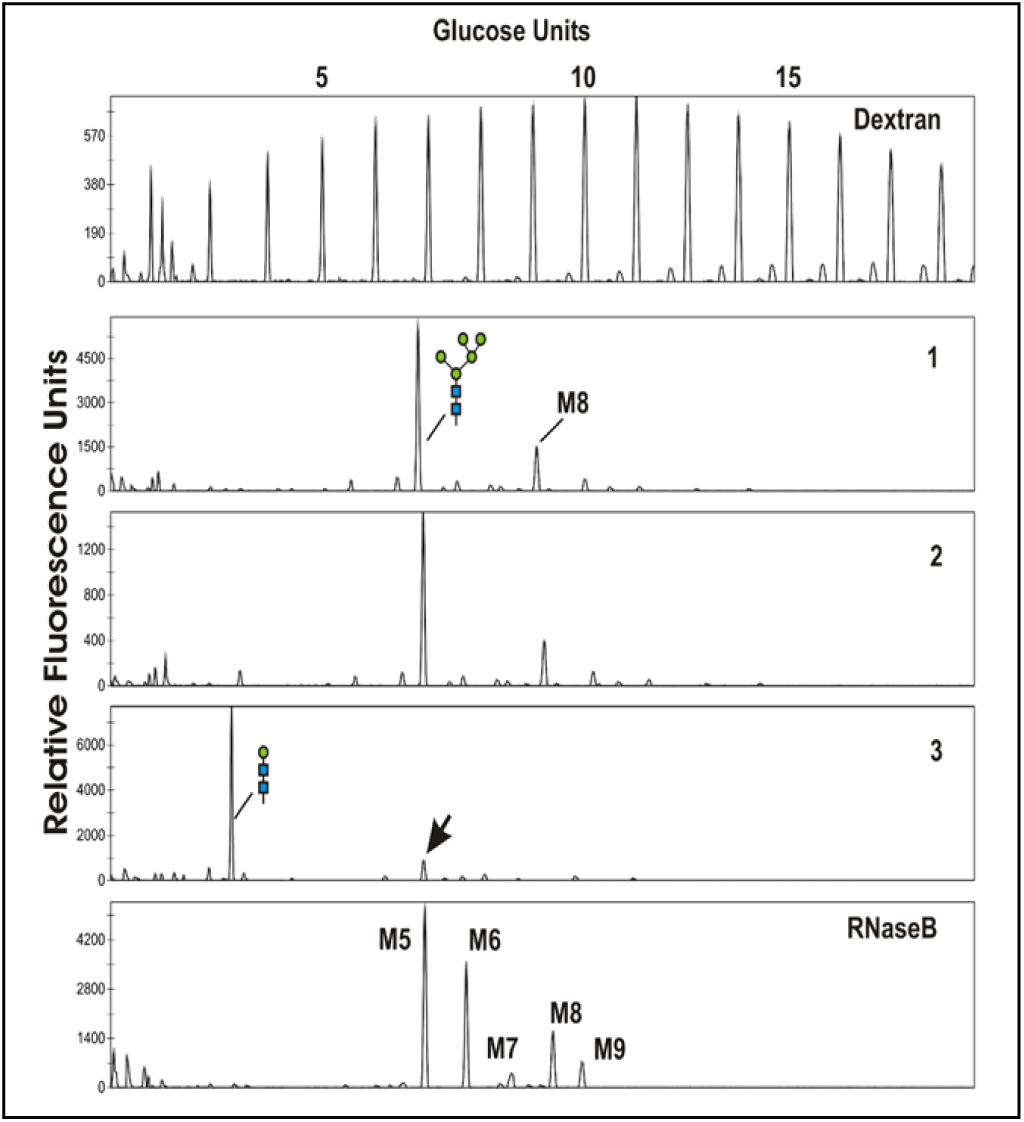
Characterization of unexpected N-glycans on GlycoSwitchM5-produced hIL-22. DSA-FACE analysis of PNGaseF-released N-glycans. Top: Dextran reference ladder; Lane 1: untreated N-glycans; Lane 2: α-1,2 mannosidase digested N-glycans; Lane 3: Jack Bean α-1,2/-3/-6-mannosidase digested N-glycans; standard N-glycan mix from RNAseB.

Given the similarities between mIL-22 and hIL-22, the N-glycans of hIL-22 were analyzed to investigate whether the glycans were structurally similar to those from mIL-22. MALDI-TOF MS analysis showed that the dominant species was Hex_5_GlcNAc_2_ (m/z 1256.35), next to two other species with m/z 1742.5 (Hex_8_GlcNAc_2_) and 1904.5 (Hex_9_GlcNAc_2_) (Supplementary Figure 3, A). MS/MS analysis of the peak at m/z 1904.5 gave a very similar fragmentation pattern as observed for the same glycan species of mIL-22 (Supplementary Figure 3, B).

We conclude that GlycoSwitchM5-hIL-22 is modified with similar off-target N-glycans as GlycoSwitchM5-mIL-22, with a different species along its biosynthesis route dominating, depending on the protein.

### Overcoming unwanted N-Glycosylation

One way of dealing with undesired N-glycans is to eliminate the N-glycan sites that carry them. To determine whether the additional N-glycan species are restricted to a specific site, three N-glycosylation site mutants of hIL-22 were constructed, each having one intact N-glycosylation site (N21, N35 or N64 of the mature hIL-22) and two Asn-to-Gln N-glycosylation site mutations. When these mutants were expressed in a GlycoSwitchM5 background, only the hIL-22 mutant with the intact N35-glycosylation site had the prominent peak around Man_8_GlcNAc_2_ on DSA-FACE (Figure 4, A), demonstrating that this modification is indeed site-specific.

**Figure 4.**
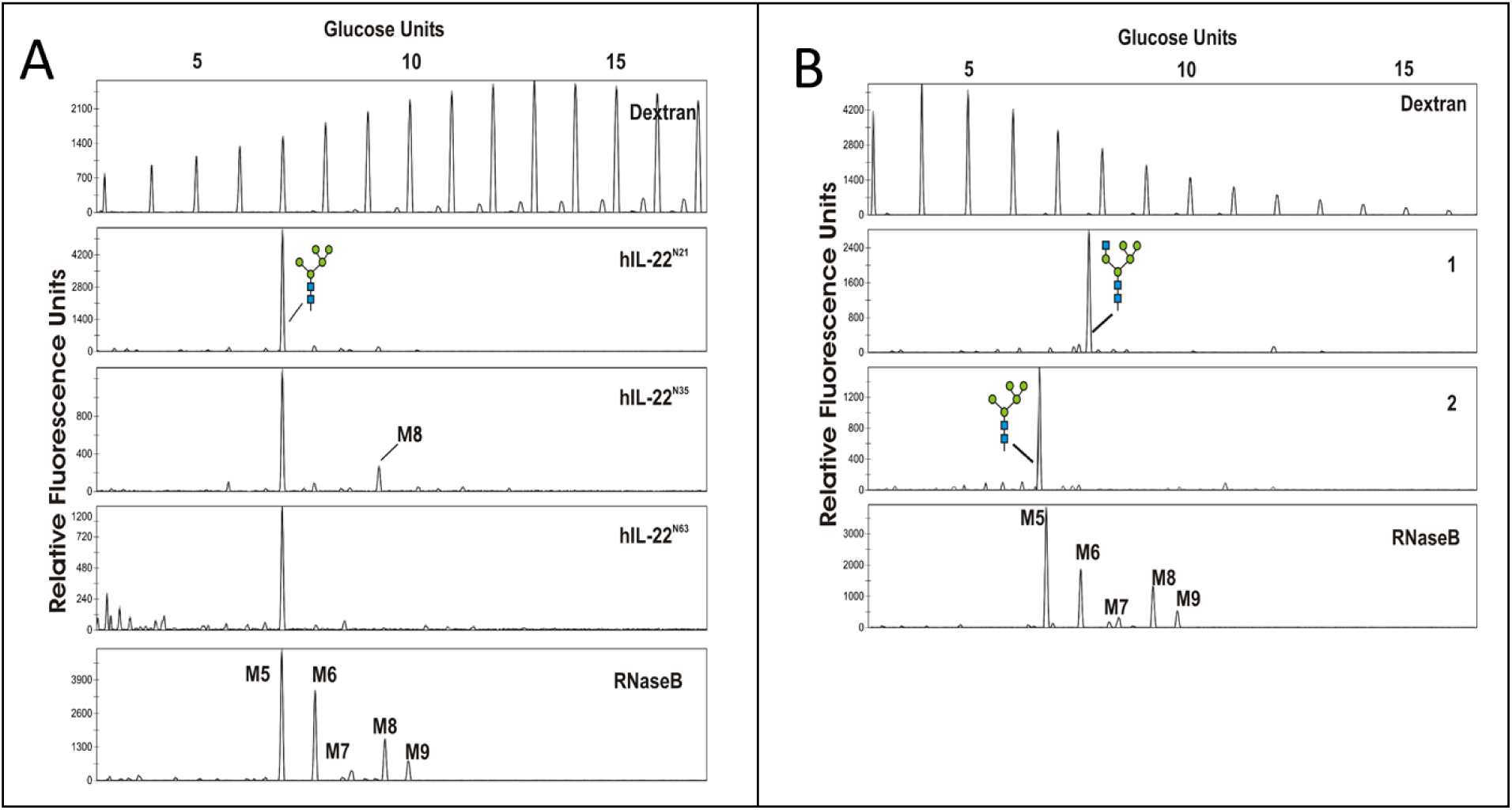
Engineering solutions to prevent the occurrence of unexpected N-glycans on GlycoSwitchM5-produced hIL-22. **A.** N-glycan analysis from three hIL-22 mutants that each contain a single N-glycosylation site either at N21(Lane 1), N35 (Lane 2) or N64 (Lane 3). **B**. N-glycan analysis from hIL-22 produced in a GlycoSwitchM5 strain that overexpresses the human N-acetylglucosaminyltransferase I (hGnT-I), either untreated (Lane 1) or treated with β-N-acetyl-hexosaminidase (Lane 2).

Another way of dealing with interference by endogenous glycosyltransferases is to outcompete them with heterologous glycosyltransferases with a similar acceptor substrate specificity. Here, the overexpression of the human N-acetylglucosaminyltransferase I (hGnT-I) was tested to see whether it is able to suppress the presence of undesired glycan species. GnT-I places a β-1,2-linked GlcNAc on the inner α-1,3 arm of the Man_5_GlcNAc_2_-core. In the strains overexpressing GnT-I, the N-glycans found on hIL-22 consisted predominantly of GlcNAcMan_5_GlcNAc_2_ (Figure 4, B; lane 1), and the peak corresponding to the glycan described above disappeared. Exoglycosidase digestion with β-N-acetyl-hexosaminidase confirmed this glycan modification by shifting the mobility back to Man_5_GlcNAc_2_ (Figure 4, B; lane 2).

## DISCUSSION

An eukaryotic host such as yeast for the expression of IL-22 can be useful since N-glycosylation can enhance the bioactivity of IL-22 (17), and IL-22 is currently being explored as a therapeutic protein (18). Therefore, the secretory expression in *P. pastoris* was explored and this showed that the glycoforms from human or mouse IL-22 had superior bioactivity compared to unglycosylated IL-22 produced in *E. coli*, demonstrating the value of the *Pichia* expression system. On the other hand, the presence of a variety of *wild-type* oligo-mannose N-glycans introduces considerable recombinant protein heterogeneity. To overcome the N-glycan associated heterogeneity, glyco-engineered *Pichia* strains are used. Previous studies have explored recombinant protein production in the GlycoSwitchM5 strain (13, 19, 20). Here the same technology was used in an attempt to express IL-22 glycoforms modified with Man_5_GlcNAc_2_ N-glycans. However, in addition to the expected Man_5_GlcNAc_2_ N-glycan, other abundant N-glycoforms were detected.

The N-glycans on glycoproteins produced by *P. pastoris* typically consist of α-linked mannose residues (21). Previously, an α-mannosidase recalcitrant Hex_9_GlcNAc_2_ N-glycan on a recombinant immunoglobulin G (IgG) from glyco-engineered *Pichia* was described by Gomathinayagam *et al*. (9). Recalcitrance to exoglycosidase digestion was reportedly due to the presence of consecutive β-linked mannose residues, which results from β-1,2-mannosyltransferase-activity that normally occurs on the outer chain of N-glycans (and O-glycans) in *wild-type P. pastoris* (7, 8). Here too, the unexpected N-glycans were recalcitrant to α-1,2 mannosidase digestion. Therefore, we initially assumed a similar structure. This was still largely in agreement with the MALDI-MS/MS data, showing a linear tetra-saccharide substituent on the termini of the Man_5_GlcNAc_2_ N-glycan. The structure identified by Gomathinayagam *et al.* appeared to be capped with a single α-1,2-mannose residue, although the authors remarkably reported that the N-glycan was recalcitrant to digestion with α-1,2 mannosidase. However, the unknown IL-22 N-glycan that we describe here was found to be capped with a glucose-residue, explaining the lack of sensitivity for α-mannosidase digestion. Glucose-residues have been reported on the outer chain of wild-type *Pichia* mannoproteins but they are linked through a phosphodiester linkage (8). A ‘capping glucose’ has been described in *Yarrowia lipolytica* as a consequence of impaired ER-processing by glucosidase-II in glyco-engineered *Δalg3* strains (22). Nevertheless, the structural data obtained by MALDI-MS/MS did not support an ER-type N-glycan.

In addition to exoglycosidase digestion and chemical linkage analysis, 1D and 2D NMR-approaches were used to elucidate more structural details of this N-glycan (6, 8, 9, 23). Although our NMR analysis did not provide full closure about what residue linked the Man_5_GlcNAc_2_ with the tetra-saccharide substitution, the combined results of all the analysis allowed us to assign a linear branch-structure consisting of Glcα-1,2-Manβ-1,2-Manβ-1,3-Glcα1-Manα1, with the latter mannose residue attached to the 3-position of one of the terminal α-mannoses of the Man_5_GlcNAc_2_ N-glycan.

The reported structure deviates considerably from what was published before (9). *Pichia* β-mannosyltransferases (BMT) are believed to add two consecutive β-1,2-linked mannose residues (6), whereas here a β-1,2- and a β-1,3-linked mannose was present, challenging this hypothesis. Only the homologous *Candida albicans* BMT1 enzyme has been biochemically studied. Analysis of *C. albicans* BMT1 showed that it transfers two consecutive β-1,2-mannose residues *in vitro*. The authors speculated that *in vivo* only a single β-mannose would be transferred by a single BMT because of competition with other endogenous mannosyltransferases present in the Golgi (24). This might be consistent with our observations where two different BMTs might act sequentially, and therefore two distinct β-mannose linkages are present in the identified structure. It is clear that more is warranted on the specificity of *Pichia* BMT enzymes in the future.

Equally puzzling is the presence of the glucose residues. The GlycoSwitchM5 strain overexpresses an HDEL-tagged α-1,2-mannosidase retaining this enzyme at the ER-Golgi boundary (19). Hence, transfer of the initiating α-1,3-linked glucose residue likely occurs at the same stage of the secretory system (or beyond) in the Golgi compartments. In the ER, the glucosyltransferases Alg6p and Alg8p take part in the dolichol-linked assembly of the precursor oligosaccharide. However, it would appear unlikely that glucosylation of the newly identified N-glycan initiates in the ER, since the addition of the linear saccharide can be outcompeted by co-expressing a Golgi localized hGnT-I. To date, no Golgi-localized α-glucosyltransferases have been described. Another hypothesis might be that the Man_5_GlcNAc_2_ N-glycan, while still in the ER, is a substrate for *Pichia* UGGT (UDP-glucose:glycoprotein glucosyltransferase) which is part of the calnexin folding cycle (25). Interaction with UGGT is not solely dependent on the N-glycan moiety but is also governed by protein-protein interactions, perhaps allowing for some promiscuity in the glycan substrate, as has been described for Trypanosomal UGGTs. However, another explanation is that yet unknown glucosyltransferases are present in the *Pichia* Golgi, especially for the most distal glucose residue which must be transferred by a transferase which acts downstream of Golgi-localized β-mannosyltransferases.

If glycoproteins are modified with undesired N-glycan modifications, mutagenesis of the affected site is by far the easiest method to deal with this, as long as the function of the protein permits it. By doing so, the α-mannosidase recalcitrant N-glycan could be traced back to the single site N35 on the mature hIL-22. Remarkably, a recent report on IL-22 produced *in planta*, showed that N35 was particularly prone to α-1,3 core fucosylation, especially when N21 was not occupied (26). Although it is unclear why only this specific N-glycosylation site is affected in hIL-22, it has been reported that the local protein surface topology as well as glycan-protein interactions or surface charge may modulate protein- or glycan-glycosyltransferase interactions and determine N-glycosylation site specific interactions (26, 27), which could also explain some differences between human and mIL-22. Nevertheless, eliminating the site at N35 seems to effectively abolish the formation of the recalcitrant N-glycan.

The tetrasaccharide modification could also be outcompeted by functional overexpression of hGnT-I. This provides for indirect evidence that the modification may have been linked to the inner α-1,3-mannose of the Man_5_GlcNAc_2_ structure as it is this residue that is modified by GnT-I. As GnT-I overexpression is the next step in the synthetic biology pathway towards human complex-type glycans (4), the formation of the off-target glycans that we describe here is strongly suppressed in these humanized strains, although it will likely not always be totally absent.

## CONCLUSION

In summary, our results show that the non-yeast native oligosaccharides resulting from *in vivo* trimming of Man_8_GlcNAc_2_ to Man_5_GlcNAc_2_ by an ER/Golgi-retained α-mannosidase serve as substrates for endogenous glycosyltransferases, creating potentially immunogenic neo-glycoforms. Although GlycoSwithM5-engineered *Pichia* strains usually modify their glycoproteins with predominantly the Man_5_GlcNAc_2_ N-glycan structure, our results demonstrate that this should be validated by careful N-glycan analysis, especially if the glyco-engineered products are to be used as biopharmaceuticals. Further innovation would be welcome to robustly remove or preclude the synthesis of these off-target high-mannose-type neo-glycans, regardless of their structure.

## MATERIALS AND METHODS

### Genetics and clone generation

*Escherichia coli* (*E. coli)* MC1061 or DH5α was used as the host strain for recombinant DNA manipulations. *E. coli* were cultivated in LB medium supplemented with the appropriate antibiotics: 100 μg/mL carbenicillin (Duchefa Biochemie) or 50 µg/mL Zeocin^®^ (Life Technologies) depending on the required selection.

The *Pichia pastoris* GS115-strain (Life technologies) was used for protein expression and is the starting strain for glyco-engineering. *P. pastoris* cells were grown in liquid YPD or solid YPD-agar and selected for with the appropriate antibiotics: 100 µg/mL Zeocin^®^ or 300 µg/mL BlasticidinS-HCl (Sigma Aldrich). Transformants of auxotrophic strains were selected on Complete Supplemented Medium (CSM) lacking Histidine (0.077% CSM-HIS, 1% D-glucose, 1.34% Yeast Nitrogen Base, 1.5% agar). Bacto yeast extract, Bacto tryptone, Bacto peptone, Bacto agar and Yeast Nitrogen Base (YNB) were purchased from Difco (Becton Dickinson). For protein expression, biomass was grown on BMGY. For induction, the cultures were switched to BMMY.

### Plasmid construction

#### pPIC9mIL22

The sequence corresponding to mature mIL-22 (UniProt accession Q9JJY9, residue 34-179) was PCR amplified using primer mIL22XhoIS and mIL22NotIAS (Supplementary Table 4) and subsequently cloned as an *Xho*I/*Not*I fragment in the similarly opened pPIC9 expression vector for *P. pastoris* resulting in the vector pPIC9mIL22, containing the mature mIL-22 coding sequence downstream of the strong methanol inducible AOX1-promoter and in-frame with the *Saccharomyces cerevisiae* α-mating factor for secretion into the culture supernatant. Selection in *P. pastoris* was carried out using the *HIS4* auxotrophic marker. Proper insert orientation was verified by restriction enzyme analysis and by Sanger sequencing using the 5’AOX1 and 3’ AOX1 primers.

#### pPIC9hIL22

The ORF corresponding to the mature form of human interleukin-22 (hIL-22, UniProt accession Q9GZX6, residue 34-179) was codon optimized for *P. pastoris* using Genscript’s proprietary algorithm and ordered synthetically as pUC57-hIL22. The hIL-22 ORF was flanked with *XhoI* and *NotI* sites to clone in-frame with the *S. cerevisiae* α mating factor in the *XhoI*/*NotI*-opened pKai and pPIC9 expression vector (28). Expression of hIL-22 in both pPIC9-hIL22 and pKai-hIL22 is under control of the AOX1 promoter. The pKai-hIL22 has a Zeocin^®^ resistance marker for selection.

### Construction of hIL22 N-glycosylation site mutants

The PCR-based Quick change site directed mutagenesis kit (Agilent) was used to remove N-glycosylation sites from the hIL-22 ORF by mutating the Asn residue of the Asn-X-Ser/Thr consensus sequence to Gln (Supplementary Table 4). By the sequential mutagenesis of the different N-glycosylation sites from pUC57-hIL22, variants were obtained that each have only a single N-glycosylation site (N21, N35 or N64). These constructs are named pUC57-IL22N21, pUC57-IL22N35 and pUC57-IL22N64 referring to the N-glycosylation site that was left intact. Each construct was sequence verified by Sanger sequencing using custom primers IL22FW and IL22RV (Supplementary Table 4). The constructs were then cloned in the pKai-expression vector as described above, yielding vectors pKai-IL22N21, pKai-IL22N35 and pKai-IL22N64. Each expression construct was sequence verified using the standard 5’ AOX1 primer and 3’AOX1 primer.

### *Pichia* transformation and strain generation

*Pichia* transformation was performed as described previously (29). All constructs were linearized to facilitate directed genomic integration. The pPIC9mIL22 vector was *Sal*I linearized in the *HIS4* marker whereas pKaiIL-22 and pPIC9-hIL22 expression constructs were *PmeI*-linearized in the AOX1-promoter prior to electroporation to the GS115 strain. For glyco-engineering of mIL-22, the GS115-strain expressing mIL-22 was transformed with the *BstB*I-linearized pGlycoSwitchM5 plasmid. The pGlycoSwitchM5 disrupts the *OCH1* gene and introduces a constitutively expressed α-1,2-mannosidase in the ER (4), generating the Man5-mIL22-strain that modifies its glycoproteins with Man_5_GlcNAc_2_. Transformation of the Man5-strain with *Sal*I-linearized pGlycoSwitchGnT-I resulted in strain GnMan5 strain that modifies its proteins predominantly with GlcNAcMan_5_GlcNAc_2_. For glyco-engineering of hIL-22, the pPIC9hIL-22 was transformed to a GlycoSwitchM5 strain, which is a GS115 strain previously transformed with the *BstB*I-linearized pGlycoSwitchM5 plasmid (4). Similarly, to generate the GnMan5-hIL-22 expression strains, the pPIC9hIL-22 expression construct was transformed to a GlycoSwitchM5 strain previously transformed with the pGlycoSwitchGnT-I (4, 19). Transformants were grown on minimal drop-out media (CSM-HIS) or YPD-agar containing the appropriate antibiotics, and were screened for IL-22 expression.

### IL-22 production in *Pichia pastoris*

Cells were grown in BMGY medium for 48 hours at 28°C (250 rpm), after which protein expression was induced by changing the growth medium to BMMY. The induction was maintained for 48 hours by spiking the cultures 2-3 times daily with 1% methanol. For clone screening, the cells were removed by centrifugation (1,500 *g* for 10 minutes) and secreted proteins were precipitated by adding 10% (v/v) of 5 mg/mL deoxycholate and 10% (v/v) of saturated trichloroacetic acid to culture supernatant. Protein was collected by centrifugation (20 minutes at 15,000x*g*, 4°C), neutralized with Tris and prepared for SDS-PAGE.

For purification of mIL-22, the culture supernatant (500 ml) was saturated with 80% ammonium sulfate. The resulting protein pellet obtained after precipitation was then resuspended in 50 mL of 25 mM Tris-acetate buffer, pH 7.8, and desalted on a Sephadex G25 XK 26/35 column (GE Healthcare). Next, the desalted protein-containing fractions were pooled and loaded on a Q sepharose FF column XK16/32 (GE Healthcare). The protein-containing fractions in the flow-through of the column were again pooled, the pH adjusted to 5.0 with 1 M HCl and loaded on a Mono S column HR10/30 (GE Healthcare), equilibrated with 25 mM acetate buffer, pH 5.0. Bound proteins were eluted with a linear gradient from 0 to 1 M NaCl in the same buffer. Finally, the mIL-22 containing fractions were pooled and loaded on a Superdex 75 XK26/60 gel-filtration column (GE Healthcare). mIL-22 protein was collected and concentrated for further analysis.

For hIL-22 purification, the ammonium sulfate precipitation pellet (as described above) was dissolved in 25mM MES pH 5.5 and filtered over a 0.22 µm bottletop filter (Millipore). The filtrate was desalted using a SephadexG25 XK26/80 column (GE Healthcare) previously equilibrated with 25mM MES pH 5.5. The desalted protein containing fractions were pooled and loaded on a Q-Sepharose XK16/32 column (GE Healthcare) equilibrated with 25mM MES pH 5.5 as running buffer. The flow through was collected and loaded on a Source 15S column. hIL-22 was eluted with a stepwise linear gradient from 0-1 M NaCl in 25 mM MES pH 5.5. The fractions containing hIL-22 were polished over a Superdex75 column (GE Healthcare) equilibrated with PBS (pH 8.0). Finally, hIL-22 containing fractions were collected and concentrated for further analysis.

### Protein deglycosylation

For DSA-FACE and MALDI-TOF MS N-glycosylation profiling, IL-22 was first denatured in 0.5% SDS, 40 mM DTT at 98°C for 10 minutes, after which the denatured proteins were supplemented with 1% of NP-40 and 50 mM Sodium Phosphate. IL-22 was then deglycosylated overnight at 37°C after adding 15.4 IUB mU of in-house produced PNGaseF.

For GC-MS and NMR analysis, a preparative N-deglycosylation method was used adapted from Aggarwal *et al.* (30). IL-22 was denatured in 0.5% SDS (w/v), 250 mM 2-mercaptoethanol, 50 mM EDTA, 250 mM potassium phosphate buffer (pH 8,6) at room temperature for 30 minutes. To each tube, four volumes of 0.2 M potassium phosphate buffer pH 8,6 was added and the reaction tubes were heated in a warm water bath for 5 minutes. The samples were deglycosylated overnight at 37°C after adding 15.4 IUB mU in-house produced PNGaseF in Triton X-100.

The released N-glycans were purified and cleaned using Carbograph Porous Graphitized Carbon (PGC) Extract-Clean columns (Alltech Associates Inc) (31, 32).

### MALDI-TOF MS

Prior to loading, the PGC columns were conditioned with 3 × 500 µL 80% acetonitrile, 0.1% TFA in MilliQ H_2_O and equilibrated with 3 × 500 µL of MilliQ H_2_O. After sample loading, the columns were washed with 3 × 500 µL of MilliQ H_2_O and free N-glycans were eluted with 3 × 500 µL of 25% acetonitrile, 0.05% TFA in MilliQ H_2_O. The eluate was pooled and evaporated to dryness.

For MALDI MS characterization, the underivatized N-glycans were spotted on a MALDI-target and mixed 1:1 with 70% MeOH, 20 µM NaCl and 20 mg/mL DHB-matrix. MALDI mass spectrometric analyses were performed on an Applied Biosystems 4800+ Proteomics Analyzer with TOF/TOF optics (Applied Biosystems, Foster City, CA). This mass spectrometer uses a 200 Hz frequency tripled Nd:YAG laser operating at a wavelength of 355 nm.

### DSA-FACE glycan analysis

APTS derivatized N-linked oligosaccharides were analyzed by DSA-FACE on a ABI 3130 capillary DNA sequencer as described previously (15). N-glycans of bovine RNaseB and a dextran ladder were included as external electrophoretic mobility standards references. Data was analyzed with the Genemapper software (Applied Biosystems). N-glycan profiles were aligned in CorelDraw 11. Exoglycosidase treatment of labeled glycans was performed with *Trichoderma reesei* α-1,2-mannosidase (produced in our laboratory, 0.33 µg per digest), Jack Bean mannosidase (Sigma, 20 mU per digest) or *Streptococcus pneumoniae* β-N-Acetylhexosaminidase (Prozyme, 4 mU per digest) overnight at 37°C in 20 mM sodium acetate (pH 5.0).

### Permethylation, mass spectrometrical analysis and linkage analysis of permethylated glycans

Permethylation of the lyophilized glycans was performed according to the procedure developed by Ciucanu and Kerek (33). The reaction was terminated by adding 1 mL of cold 10% (v/v) acetic acid followed by three extractions with 500 µL of chloroform. The pooled chloroform phases were then washed eight times with water. The chloroform phase containing the methylated derivatives was finally dried under a stream of nitrogen and the extracted products were further purified on a C18 Sep-Pak (24). The C18 Sep-Pak was sequentially conditioned with methanol (5 mL), and water (10 mL). The derivatized glycans dissolved in methanol were applied on the cartridge, washed with 3 x 5 mL water, 2 mL of 10% (v/v) acetonitrile in water and eluted with 3 mL of 80% (v/v) acetonitrile in water. The acetonitrile was evaporated under a stream of nitrogen and the samples were freeze-dried. MALDI-TOF-MS and linkage analyses of permethylated glycans were performed as described elsewhere (34).

### NMR analysis

All NMR spectra were measured on a Bruker Avance II NMR spectrometer operating at frequencies of 700.13 MHz and 176.05 MHz for ^1^H and ^13^C respectively. All measurements were performed using a 1 mm TXI ^1^H-^13^C/^15^N Z-gradient probe for maximal signal intensity. For sample preparation, the lyophilized N-glycans were dissolved in 10 µl of 99.9% D_2_O. 2,2-dimethyl-2-silapentane-5-sulfonate (DSS) was used as an internal chemical shift standard and 1 µl of NaN_3_ was added to prevent microbial growth. The mixture was then transferred to a 1 mm capillary tube (Bruker). All measurements were performed at 50°C to minimize overlap between the residual water signal and those of the N-glycan.

The 2D COSY (Correlation Spectroscopy), TOCSY (Total Correlation Spectroscopy), ^1^H-{^13^C}-gHSQC (Heteronuclear Single Bond Quantum Coherence) and ^1^H-{^13^C}-gHMBC (Heteronuclear Multiple Bond Coherence) were measured using standard sequences available in the Bruker pulse programme library (cosyprqf, mlevphpr, noesyphpr, hsqcedetgpsisp2, hmbcgplpndqf). The mixing time for TOCSY and NOESY was set to 100 and 200 ms, respectively. The homonuclear 2D experiments were performed with 512 increments, accumulated from 32 to 64 scans of 2048 data points each. The NMR relaxation time was 1.27 to 1.50 s and the ^1^H spectral width was 11.30 ppm. Data were multiplied using a squared sine window (COSY) or a squared cosine window (TOCSY and NOESY) before zero-filling to a 2048 by 2048 real data matrix. A second order polynomial baseline correction was performed on both the F1 and F2 dimension. For heteronuclear 2D experiments, a ^13^C spectral width of 160 ppm (gHSQC) and 220 ppm (gHMBC) was used, and 8 to 64 scans were accumulated for each of the 512 increments. Data were multiplied with a squared sine window, followed by zero filling as before, and Fourier Transformation followed by second order baseline correction, when necessary.

### Interleukin-22 bio-activity assay

The biological activity of both murine and human Man_5_GlcNAc_2_-IL-22 was determined using the mouse pro-B cell line Ba/F3m22R (Ba/F3 cells expressing the mouse IL-22 receptor (IL22R1) as described in (35); Ba/F3m22R cells were seeded in 96-well plates at a concentration of 3000 cells per well in the presence of dexamethasone. Three days after addition of serial sample dilutions, cell growth was measured by colorimetric determination of hexosaminidase levels according to Landegren (36). The EC_50_ was determined based on the dose-response curve and the specific activity was calculated as A(U/mg)=(1/EC_50_)*10^6^. Mouse IL-22 expressed in *E. coli* (ECmIL-22) was purified as described previously (37) and was used as control.

## ACKNOWLEDGEMENTS

B.L., PP. J. and C. DW. held a fellowship of the Institute for the Advancement of Scientific and Technological Research in Industry (IWT). This work was in part funded by grant no. G.0.541.08.N.10 of FWO-Vlaanderen and an ERC Consolidator grant (GlycoTarget) to N.C.. We thank Ingeborg Stals for HPAEC-PAD analysis. W.M. was supported by the Centre National de la Recherche (Unité Mixte de Recherche CNRS/USTL 8576), and the Ministère de la Recherche et de l’Enseignement Supérieur. The Mass Spectrometry facility used in this study was funded by the European Community (FEDER), the Région Nord-Pas de Calais (France) and the Université des Sciences et Technologies de Lille.

We dedicate this paper to the memory of Dr. Bart Samyn who sadly passed away during the course of these studies.

## AUTHOR CONTRIBUTIONS

B.L. wrote the manuscript, designed and performed the experiments concerning hIL-22. PP.J. engineered the mIL-22 strains, purified mIL-22 and performed initial N-glycan analysis, K.G. and J.M. performed the NMR analysis, C. DW aided in the mutagenesis and expression of hIL-22, P.A. performed N-glycan analysis, W.M. performed analysis on permethylated N-glycans, B.S. performed the MS analysis, J-C. R. coordinated the IL-22 bioactivity assays. R.C. and N.C. designed experiments, interpreted, and supervised the work. S.D. and N.C. co-wrote the manuscript.

