## Supplementary Information for "Off-target glycans encountered along the synthetic biology route towards humanized N-glycans in *Pichia pastoris*"

### Supplementary Figures

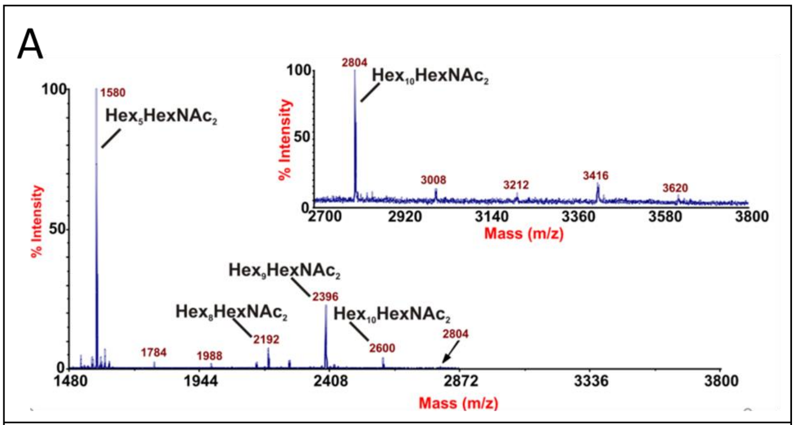


B

| Ion at m/z | Abundance % | N-glycan |
|---|---|---|
| 1580 | 72.6 | $Hex_5HexNAc_2$ |
| 1784 | 1.8 | $Hex_6HexNAc_2$ |
| 1988 | 1.3 | $Hex_7HexNAc_2$ |
| 2192 | 5.0 | $Hex_8HexNAc_2$ |
| 2396 | 16.0 | $Hex_9HexNAc_2$ |
| 2600 | 3.0 | $Hex_{10}HexNAc_2$ |
| 2804 | 0.4 | $Hex_{11}HexNAc_2$ |

**S-Figure 1. MALDI-TOF MS analysis of the permethylated N-glycan pool. A.** MS spectrum that was used to calculate the abundance of each N-glycan species. Inset: zoom of the range m/z 2700 to 3800. **B.** The abundance of the different N-glycans present on GlycoSwitchM5-produced mIL-22.

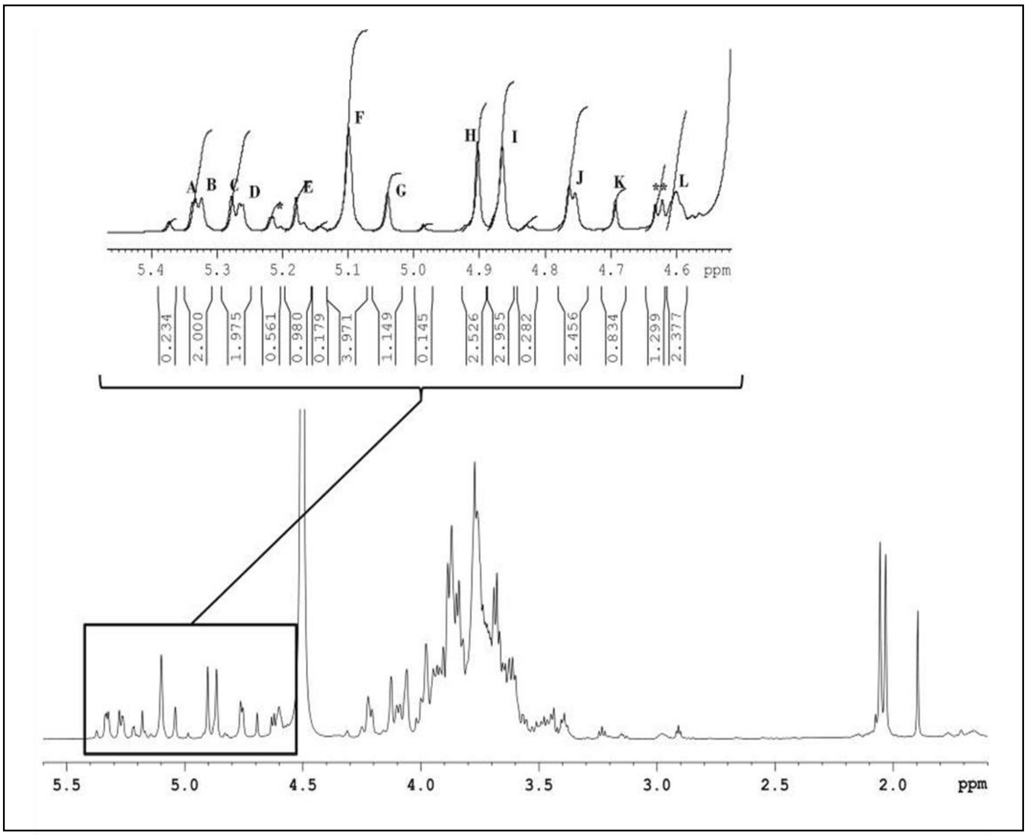


**S-Figure 2. 700 MHz 1D $^1$H-NMR spectrum of GlycoSwitchM5-produced mIL-22 N-glycans.** The anomeric region was enlarged (box) to enable comparison of the relative intensities between signals (A-L, * and **) from the N-glycan sample containing $Man_5GlcNAc_2$ and the unknown N-glycan. Anomeric signals marked with * and ** were identified as monosaccharide contaminants trough diffusion analysis. Signals that were not annotated were also eliminated based on their low relative intensities. After removal of all potential contaminants, signal A-L were identified as true components of the unknown N-glycan.

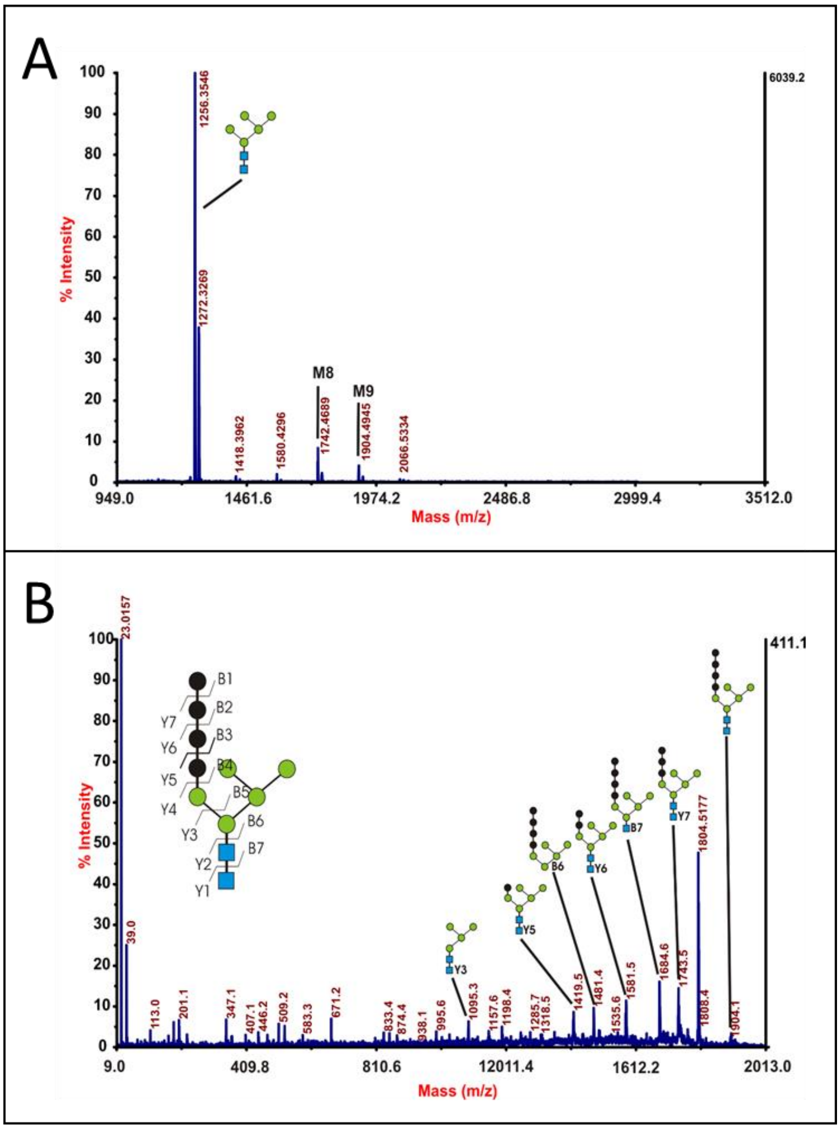


**S-Figure 3. MALDI-TOF MS analysis of the released N-glycans from GlycoSwitchM5-produced hIL-22. A.** MS spectrum of the released N-Glycans. **B.** MS/MS fragmentation of the $Hex_9GlcNAc_2$ species (m/z 1905.6).

### Supplementary Tables

**S-Table 1. GC-MS linkage-analysis on permethylated and acid hydrolyzed N-glycans.** Permethylated glycans were hydrolyzed, reduced, acetylated and analyzed by GC-MS. The retention times and the characteristic fragment ions used to assign an identity to the hydrolyzed monosaccharides is shown together with the relative abundance of each.

| Retention time (min) | Characteristic fragment ions | Assignment | Abundance (% total) |
|---|---|---|---|
| 24.26 | 102, 118, 129, 145, 161, 162, 205 | Terminal glucose | 5.4 |
| 24.43 | 102, 118, 129, 145, 161, 162, 205 | Terminal mannose | 27.2 |
| 28.56 | 129, 130, 161, 190 | 2-linked mannose | 15.2 |
| 29.00 | 101, 118, 161, 234 | 3-linked glucose | 8.4 |
| 29.43 | 101, 118, 161, 234 | 3-linked mannose | 5.7 |
| 30.24 | 102, 118, 129, 162, 189, 233 | 6-linked mannose | 1.1 |
| 33.49 | 129, 130, 189, 190 | 2,6-linked mannose | 1.1 |
| 34.24 | 118, 129, 189, 234 | 3,6-linked mannose | 22.8 |
| 38.46 | 117, 159, 233 | 4-linked GlcNAc | 10.3 |

**S-Table 2. Diffusion coefficient of the anomeric signals.** The diffusion coefficient of the anomeric signals in the sample was determined to establish whether all signals are part of the same structure. In the latter case, the coefficient should be similar for every signal. Higher coefficients point towards free monosaccharides (contaminants) in the sample (* and **).

| | D (*$10^{-10}$ $m^2/s$) |
|---|---|
| A | 3.12 |
| B | 3.02 |
| C | 2.86 |
| D | 2.96 |
| E | 3.55 |
| F | 3.56 |
| G | 2.81 |
| H | 3.15 |
| I | 3.31 |
| J | 3.85 |
| K | 2.77 |
| L | 3.81 |
| * | 8.88 |
| ** | 10.57 |

**S-Table 3. HSQC $^{13}$C data of the N-glycan sample and their GS.** The GS is obtained through comparison of the $^{13}$C data with the data of the individual monosaccharides. Values indicated in bold indicate the presence of a glycosidic linkage. For each monosaccharide, values on the same line represent the $^{13}$C chemical shift of the respective C-atoms of the monosaccharide ring. The value beneath indicates the GS. Lettering is the same as in the $^{1}$H- and TOCSY spectrum. The inset glycan-scheme reflects the lettering in the structure of the N-glycan.

| Kernstructuur | Trace | C-1 | C-2 | C-3 | Extra | Trace | C-1 | C-2 | C-3 |
|---|---|---|---|---|---|---|---|---|---|
| α-D-Man-(1→ | **F** | 104.99 | 72.88 | 73.18 | →3)-α-D-Glc-(1→ | **A** | 101.34 | 74.56 | 83.35 |
| | | **9.49** | 0.58 | 1.28 | | | **7.74** | 1.36 | **8.85** |
| →3,6)-β-D-Man-(1→ | **J** | 103.00 | 72.79 | 83.39 | →2)-β-D-Man-(1→ | **B** | 103.43 | 81.16 | overlap |
| | | **7.80** | -0.01 | **8.59** | | | **8.23** | **8.36** | |
| →3,6)-α-D-Man-(1→ | **H** | 102.16 | 72.78 | 78.65 | →2)-β-D-Man-(1→ | **C** | 103.25 | 81.16 | overlap |
| | | **6.66** | 0.48 | **6.75** | | | **8.05** | **8.36** | |
| α-D-Man-(1→ | **I** | 102.57 | 72.23 | 73.18 | α-D-Glc-(1→ | **D** | 103.26 | 69.64 | overlap |
| | | **7.07** | -0.07 | 1.28 | | | **9.66** | -3.56 | |
| α-D-Man-(1→ | **F** | 104.99 | 72.88 | 73.18 | →3)-α-D-Man-(1→ | **G** | 104.78 | 72.82 | 81.30 |
| | | **9.49** | 0.58 | 1.28 | | | **9.28** | 0.52 | **9.40** |
| →4)-β-D-GlcNAc-(1→ | **L** | 104.17 | 57.85 | overlap | | | | | |
| | | **6.77** | -18.05 | | | | | | |
| →4)-α-D-GlcNAc | **E** | 93.22 | 56.44 | overlap | | | | | |
| | | -0.38 | -16.76 | | | | | | |
| →4)-β-D-GlcNAc | **K** | 97.53 | 58.97 | overlap | | | | | |
| | | 0.13 | -16.93 | | | | | | |

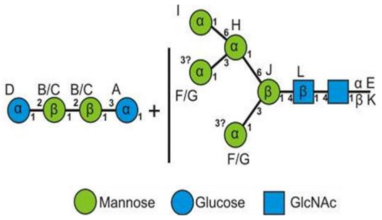

**S-Table 4. Primers used in this study**

| | |
|---|---|
| mIL22XhoIS | 5’-CTAGCTCGAGAAAAGAGAGGCTGAAGCCCTGCCCGTCAACACCCGGTGC-3’ |
| mIL22NotIAS | 5’-AGTTTAGCGGCCGCTCAGACGCAAGCATTTCTCAGAGAC-3’ |
| a115c_117a_FW | 5’-TGGCTAAAGCTTCTCTGGCTGATCAAAACACCGATGTTCG-3’ |
| a115c_117a_RV | 5’-CGAACATCGGTGTTTTGATCAGCCAGAGAAGCTTCTTTAGCCA-3’ |
| a202c_c204a_FW | 5’-GAAACAGGTTCTGCAATTTACCCTGGAAGAAGTTCTGTTTCCACAG-3’ |
| a202c_c204a_RV | 5’-CCAGGGTAAATTGCAGAACCTGTTTCATCAGATAACAACGTTC-3’ |
| IL22IValtFW | 5’-GCCATACATCACCCAACGTACCTTTATGCTGGC-3’ |
| IL22IVRV | 5’-CAGCATAAAGGTACGTTGGGTGATG-3’ |
| IL22FW | 5’-CTCGAGAAACGTGCTCCAATTTC-3’ |
| IL22RV | 5’-GCGGCCGCACTAGTCTATTAC-3’ |
| 5’AOX1 | 5’-GACTGGTTCCAATTGACAAGC-3’ |
| 3’AOX1 | 5’-CAAATGGCATTCTGACATCC-3’ |
